# Reorganisation of circadian activity and the pacemaker circuit under novel light regimes

**DOI:** 10.1101/2024.05.07.592876

**Authors:** Pragya Sharma, Vasu Sheeba

**Affiliations:** Neuroscience Unit, Jawaharlal Centre for Advanced Scientific Research, Bangalore, India

## Abstract

Many environmental features are cyclic, with predictable daily and yearly changes which vary across latitudes. Organisms cope with such change using internal timekeepers or circadian clocks which have evolved remarkable flexibility. This flexibility is evident in the waveforms of behavioural and underlying molecular rhythms. In today’s world, many ecosystems experience artificial light at night, leading to unusual photoperiodic conditions. Additionally, occupational demands expose many humans to unconventional light cycles. Yet, practical means of manipulating activity waveforms for beneficial purposes are lacking. This requires an understanding of principles and factors governing waveform plasticity of activity rhythms. Even though waveform plasticity remains underexplored, few recent studies have used novel light regimes, inspired by shift work schedules, with alternating bright light and dim light (LD^im^LD^im^) to manipulate the activity waveform of nocturnal rodents. We undertook this study to understand what aspects of light regimes contribute to waveform flexibility and how the underlying neuronal circuitry regulates the behaviour by subjecting *Drosophila melanogaster* to novel light regimes. Using a range of LD^im^LD^im^ regimes, we found that dim scotopic illumination of specific durations induces activity bifurcation in fruit flies, similar to mammals. Thus, we suggest evolutionarily conserved effects of features of the light regime on waveform plasticity. Further, we demonstrate that the circadian photoreceptor CRYPTOCHROME is necessary for activity bifurcation. We also find evidence for circadian reorganisation of the pacemaker circuit wherein the ‘evening’ neurons regulate the timing of both bouts of activity under novel light regimes. Thus, such light regimes can be explored further to understand the dynamics and coupling within the circadian circuit. The conserved effects of specific features of the light regime open up the possibility of designing other regimes to test their physiological impact and leverage them for waveform manipulation to minimise the ill effects of unusual light regimes.

**Author Summary:** It is thought that the appropriate timing of physiological and behavioural rhythms of organisms with respect to the environmental cycle confers an adaptive advantage. Endogenous timekeepers or circadian clocks regulate such rhythms. To optimally time biological rhythms, its waveform must be plastic and respond to changes in external cycles. Changes in external cycles may be natural, as seen across latitudes or seasons, or anthropogenic, such as artificial light induced changes in photoperiod or shiftwork driven novel light/dark cycles. Previous studies using a nocturnal rodent model showed that novel light regimes (LD^im^LD^im^) caused locomotor activity to bifurcate such that mice showed two bouts of activity restricted to the dimly lit phases.

Here, we first demonstrate that conserved features of the light regime - dim scotopic illumination of specific light durations induce activity bifurcation in the fly model. We leverage the genetic toolkit of the Drosophila model to also show evidence for the reorganisation of the circadian pacemaker neuronal network upon exposure to novel light regimes. Our findings indicate that conserved effects of specific features of the environmental regimes can be exploited to design light regimes that ease the waveform into synchronising with challenging conditions such as during shift work, jetlag, and photoperiodic changes.

## Introduction

Cyclic geophysical factors have shaped the evolution of organisms across millennia. Periodic changes in light and temperature across a daily and seasonal scale are among the most apparent abiotic rhythmic phenomena. They are associated with rhythmic physiological and behavioural processes in almost all life forms. It is thought that organisms have evolved internal mechanisms called circadian clocks to synchronise biological processes to environmental cycles appropriately. However, cyclic environmental inputs are variable across seasons and latitudes. Such variation demands that the circadian clock and its output rhythms be flexible to generate appropriate changes in the behavioural or physiological rhythm waveform. This lability in clock function is likely to be adaptive. The response of activity waveform to photoperiodic changes, jetlag, shift work or even daylight savings indicates that the circadian system strikes a delicate balance between rigidity and plasticity. This would allow the system to be labile enough to respond to changes but also maintain robust timing in the face of minor environmental fluctuations. Despite detailed mechanistic insights into circadian clock functioning, an understanding of the practical means by which rhythm waveforms can be efficiently manipulated is lacking. This would require a better understanding of the principles and factors governing waveform plasticity of biological rhythms.

Among all the behavioural rhythms exhibited by metazoans, daily rhythmicity in locomotion has garnered significant interest due to its apparent utility in providing insights into waveform regulation. While the waveform of locomotor activity is considered a reasonably close reflection of the state of the underlying circadian machinery, factors such as startle responses or timing of food availability are also known to influence it [1–3]. Thus, it is crucial to understand clock and non-clock factors regulating activity waveform. Even though waveform plasticity has received limited attention, a few studies have used novel light regimes to probe the waveform plasticity of locomotor activity in rodents [4,5]. Through these, a unique phenomenon has been described in Siberian and Syrian hamsters exposed to light regimes with two alternating bright and dim light phases (LD^im^LD^im^) within 24 hours. Under this regime, the activity waveform comprises two segregated bouts restricted to the dim light phases. This phenomenon is distinct from the activity pattern of the same organisms under standard LD 12:12 conditions and is termed ‘bifurcation’ or behavioural decoupling. Dim scotopic illumination was found to be critical for activity bifurcation since a similarly timed regime with darkness in the scotophases (LD^ark^LD^ark^) produced only a single bout of activity across 24 hours [6].

At the cellular level, such bifurcation of locomotor activity was correlated with antiphasic oscillations of both PER1 and BMAL1 between the core and the shell of the SCN in mice [7,8]. It is thought that such behavioural plasticity reflects the inherent rigidity and plasticity of the circadian system and depends on the ability of dim scotopic illumination to influence coupling within the circadian circuit [6,9,10]. Through this series of studies using the model system of nocturnal rodents, for which the neuronal network dynamics are known to a considerable extent, the authors unravel factors contributing to waveform flexibility, namely, dim scotopic illumination, duration of light phases, and the reorganisation of the neuronal network. Considering that the above studies are limited to nocturnal rodents, it is impossible to generalise how circadian systems, behaviours or physiological processes respond to such environmental challenges unless studied across taxa, especially those that otherwise occupy different ecological and temporal niches.

In a previous study by our group, *Drosophila melanogaster* flies were subjected to a light regime with alternating light and dark phases (LD^ark^LD^ark^) to probe the flexibility of activity waveform and oscillator or network interactions [11]. There, we demonstrated that, similar to rodents, fly activity also does not bifurcate under LD^ark^LD^ark^. Instead, the waveform is similar to that under a long photoperiod, albeit with a less prominent evening peak. This observation and the fact that dim scotopic illumination was crucial for activity bifurcation in rodents led to the following questions: a) Would dim scotopic illumination induce bifurcation of activity waveform in flies similar to nocturnal rodents? and b) If so, what aspects of the light regime contribute to it – duration of the light phases, wavelength, and/or intensity? We reasoned that our studies using the fly could unravel common principles governing circadian oscillator decoupling and waveform plasticity. Additionally, we leveraged the genetic toolkit of *D. melanogaster* to examine the role of photoreception, circadian clock genes and the cellular components which house the oscillator/s regulating the activity pattern under novel light regimes.

We exposed *Drosophila melanogaster* to a series of light regimes based on those previously described for rodents. Alternating bright and dim light phases (LD^im^LD^im^) were provided spread across 24 hours. This was repeated over 5-7 cycles, and activity waveforms were characterised. We find that *D. melanogaster* undergoes activity bifurcation under certain combinations of durations of the light and dimly lit phases. Additionally, we reveal a role for the Drosophila circadian photoreceptor CRYPTOCHROME and canonical circadian clock genes in determining the waveform under this regime. Our study also uncovers unique features of the fly circadian network, which suggest that specific subsets previously described as ‘evening’ cells can modulate the phasing of both bouts when activity is bifurcated under this novel regime.

## Results

### Locomotor activity bifurcation in *Drosophila melanogaster*

In rodents, LD^ark^LD^ark^ does not induce bifurcation of activity. However, bifurcation was observed upon introducing dim scotopic illumination to this regime [4]. Previous work from our group characterised activity pattern in *Drosophila melanogaster* under LD^ark^LD^ark^ [11]. Similar to rodents, activity bifurcation was not observed in fruit flies under this light regime, which led to the question of whether dim scotopic illumination was required for a similar effect on the activity waveform in flies, i.e. bifurcation. *Drosophila melanogaster* has been previously shown to exhibit heightened activity under moon-lit conditions when subjected to light: moonlight 12:12 regimes [12]. Thus, we first assayed the activity of flies under LD^im^ 12:12 (with 70 lux: 1 lux), wherein we observed a large evening bout, a reduced morning bout, and increased nocturnal activity (S1 Fig). Following this, we recorded activity under LD^im^LD^im^ to test if introducing dim light during the scotophases would induce and maintain stable bifurcation, as in rodents [4]. The flies were under LD 12:12 for the first few days. Then they were shifted to a light regime with alternating bright light (70 lux) and dim light (1 lux), LD^im^LD^im^ 5:7:5:7. Interestingly, we observed bifurcation - the activity was restricted to and symmetrically distributed between the two dim light phases, with very little activity during the bright light phases (Fig 1). This is unlike LD12:12 or LD^im^ 12:12 (S1 Fig). To verify that such bifurcation and dim light preference is not unique to a particular strain, we assayed activity/rest under the LD^im^LD^im^ regime for flies of three different strains - one from a well-studied laboratory population with a considerable standing genetic variation called *control*, described in a previous study [11] and referred to as GCs here, and common inbred laboratory ‘wild type’ strains of *Canton-S* and *w^1118^* (Fig 1A). The batch actogram of the GC flies (Fig 1B) shows the activity distribution under LD 12:12 followed by transfer to LD^im^LD^im^ 5:7:5:7, under which they appear to bifurcate their activity. We used bifurcation index to quantify the pattern observed [5] (for details, see materials and methods). This index will only assume a value of 1 if all the activity is symmetrically distributed between the two scotophases (dim light phases) with no activity during the photophases. A value significantly higher than 0.5 indicates that most activity is restricted to the scotophases and symmetrically distributed, thus indicative of a bifurcated activity pattern. For all three strains assayed under this regime, the values of the bifurcation index were significantly higher than 0.5 by one-sample *t*-tests (Fig 1C). Also, a sudden startle or masked response was observed at the bright-to-dim light transition. These observations across strains show that the effect of exposure to LD^im^LD^im^ on activity waveform is similar between rodents and the fruit fly, i.e. bifurcation or behavioural decoupling.

**Fig 1.**
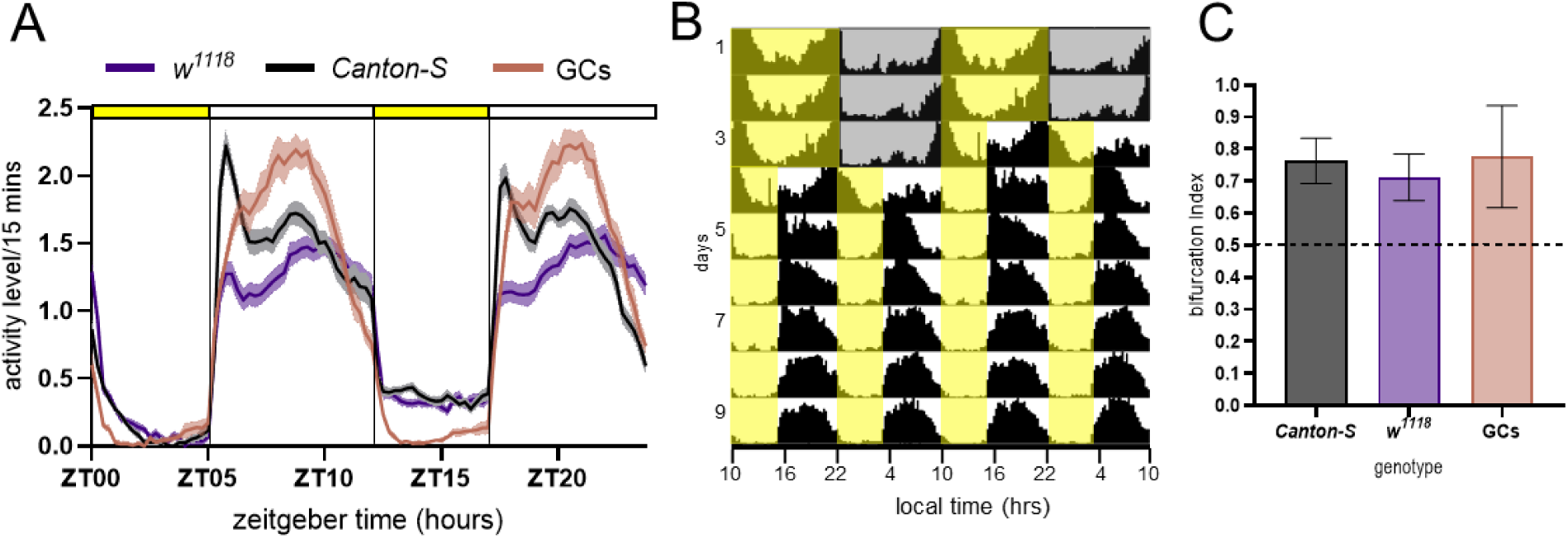
Activity bifurcation in *Drosophila melanogaster*. (A) Activity profiles averaged over cycles and across individuals of the populations along with the inbred lines of *w^1118^*and *Canton-S* under LD^im^LD^im^ 5:7:5:7. Error bands are SEM. The yellow and white bars at the top indicate bright (70 lux) and dim light (1 lux), respectively. (B) Batch actogram of flies from outbred populations (GCs) exposed to LD 12:12 with 70 lux light intensity followed by LD^im^LD^im^ 5:7:5:7 with 70 lux of bright light alternating with 1 lux of dim light. Yellow shading indicates 70 lux, while grey shading is for complete darkness. In the following cycles of LD^im^LD^im^, yellow indicates 70 lux, while non-shaded zones indicate 1 lux. (C) Bifurcation index of *Canton-S*, *w^1118^* and GC fly strains (for each strain *n* > 50). The bifurcation index (BI) was averaged over cycles for each individual and then across individuals. BI is significantly higher than 0.5 by one-sample *t*-test for all three strains (*p* < 0.0001). Error bars are SD.

### Activity bifurcation in *D. melanogaster* occurs under unique combinations of light: dim durations

To assess the properties of the light regime influencing activity bifurcation, we screened a series of LD^im^LD^im^ regimes since such a peculiar waveform where activity is highly restricted to the scotophases may be produced only by a particular sequence and duration of bright and dim light cycles. We tested the following hypotheses –

**Hypothesis 1**: Symmetrical light regimes facilitate waveform bifurcation. We reasoned that LD^im^LD^im^ 5:7:5:7 is a symmetrical regime wherein there are two consecutive short light-cycles/T-cycles in 24 hours and, thus, can be considered entrainment to 12-hour days (LD^im^ 5:7). We examined whether the activity could entrain with a bifurcated pattern to other combinations of T-cycles which we termed ‘asymmetrical’ since the alternate T-cycles were not similar. Under the regime LD^im^LD^im^ 5:5:5:9, we observed that activity bifurcated into the two scotophases (Fig 2B). Hence, we negated this hypothesis and concluded that the activity also bifurcates under asymmetrical light regimes, indicating synchronisation to 24-hour cycles.
**Hypothesis 2**: Bifurcation occurs when the total duration of dim light is greater than that of bright light. The total duration of dim versus the total duration of bright light could also be a factor that induces bifurcation. This would be akin to the observation of specific waveforms only under certain photoperiods; for example, under a long photoperiod, a diminished morning bout and a prominent evening bout of activity are observed [13]. In both regimes described thus far, the total duration of dim light total was greater than that of bright light (14 hours versus 12 hours). Therefore, we next imposed LD^im^LD^im^ 5:5:7:7, which has equal durations of total bright and dim light (12 hours each). The observation of bifurcated activity under this regime indicates that the total duration of dim light need not be greater than that of bright light for such a pattern to be observed (Fig 2C). So far in our study, the regimes used had a total dim light duration equal to or greater than the total bright light. To test the necessity of such a ratio, we provided LD^im^LD^im^ 7:5:7:5 (10 hours of dim light as opposed to 14 hours of bright light). A bifurcated activity waveform was also observed under this regime (S1 Fig).
**Hypothesis 3**: A critical minimum scotophase duration is required for bifurcation of activity. We exposed the flies to LD^im^LD^im^ 5:3:7:9, where the total duration of bright and dim light was equal (12 hours). Here, the durations of each of the photophases are the same as the previous regime (LD^im^LD^im^ 5:5:7:7). However, in this regime, we introduced a shorter duration (3 hours) for the first scotophase. We found that the activity did not bifurcate (Fig 2D). We also exposed *Canton-S* flies to LD^im^LD^im^ 5:7:5:7 and then transferred them to 5:3:7:9 to confirm and longitudinally visualise the effect of the regime on the waveform (Fig 2E). Thus, we find that there is a threshold duration of the scotophase (> 3 hours) that facilitates activity bifurcation.

**Fig 2.**
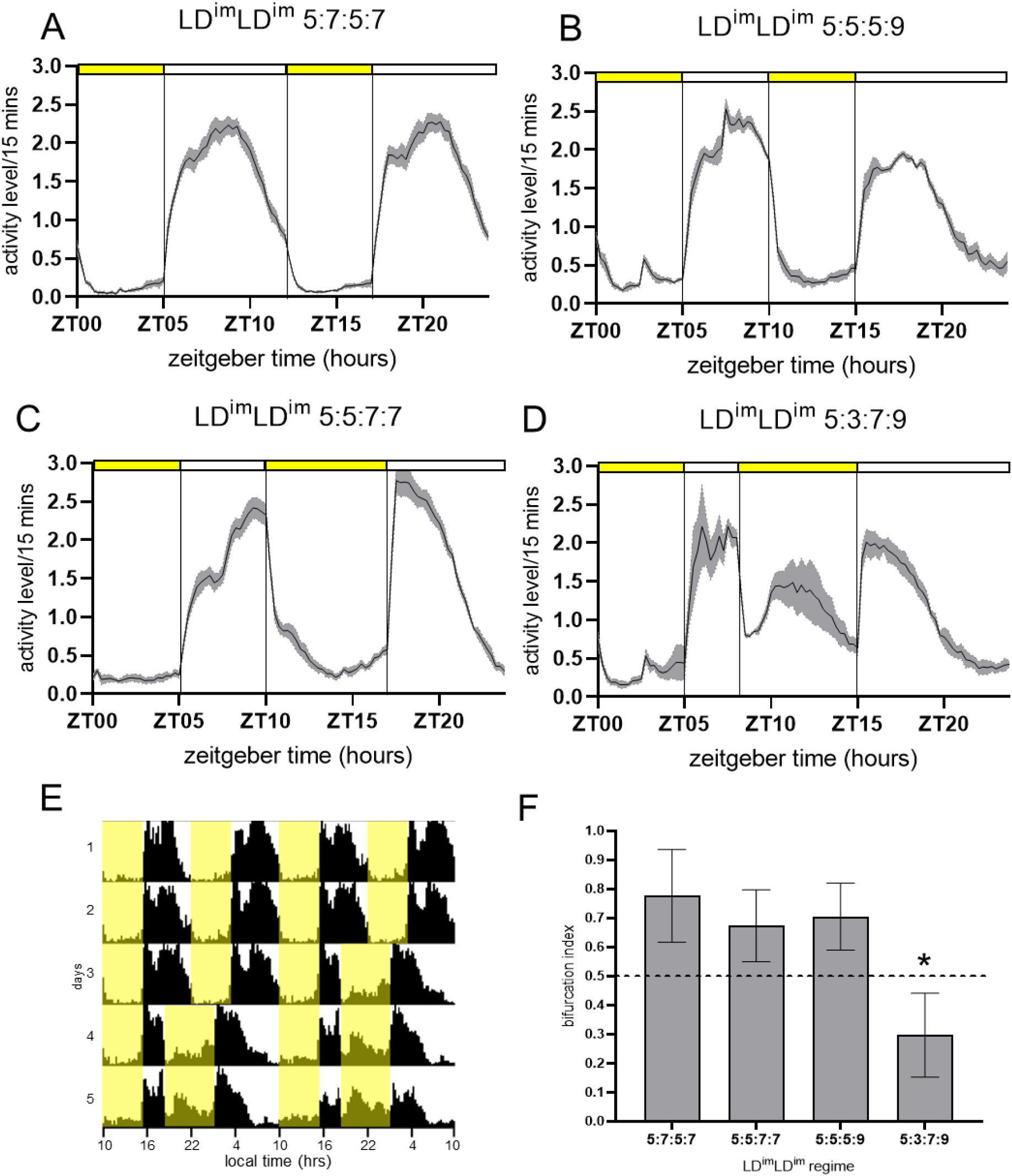
Activity bifurcation depends on scotophase duration. (A-D) Activity profiles of flies from GC populations of *Drosophila melanogaster* under four different light regimes. The regime for each profile is indicated at the top, with the yellow and white bars indicating the bright and dim light durations, respectively. Profiles have been averaged over cycles and across individuals; error bands are SEM. (E) Batch actogram of *Canton-S* flies exposed to LD^im^LD^im^, yellow indicates 70 lux, while non-shaded zones indicate 1 lux. The flies were first exposed to LD^im^LD^im^ 5:7:5:7 for three cycles and then to LD^im^LD^im^ 5:3:7:9 for the subsequent three cycles to visualise the change in activity waveform from a bifurcated one to a non-bifurcated one as a function of the light regime. (F) Bifurcation index of flies under each of the four LD^im^LD^im^ regimes. The asterisk indicates that the value was significantly lower than 0.5 by a one-sample *t*-test. In contrast, the absence of an asterisk indicates that BI was significantly higher than 0.5. For each of the four comparisons, *p* < 0.0001, *n* > 84, and error bars are SD.

Bifurcation indices under the above light regimes were estimated, and the value was significantly lower than 0.5 only for LD^im^LD^im^ 5:3:7:9 (Fig 2F). Using this series of regimes, we found that there is a requirement of a critical minimum duration of the scotophase and that activity bifurcation can also occur under asymmetrical regimes and even under regimes with a total dim light duration less than the bright light duration. In the case of nocturnal rodents as well, specific photoperiodic requirements were crucial for stable induction of behavioural decoupling [14]. Thus, our results highlight that along with dim scotopic illumination, threshold durations of scotophases and photophases could also be a conserved factor determining the induction of such behavioural decoupling.

### Bifurcated activity restricted to dim light is mediated by *cryptochrome* in the circadian clock neurons

Our experiments indicated that dim scotopic illumination of a specific duration is crucial for activity bifurcation. Thus, factors involved in dim light detection would play a significant role in regulating activity waveforms under these conditions. Fruit flies have three major pathways for photoreception: HB eyelets, compound eyes and the cell-autonomous photoreceptor CRYPTOCHROME (CRY) expressed in subsets of the circadian pacemaker neurons. CRY is known to sense lower light intensities, compound eyes moderate light intensities, while HB eyelets sense higher light intensities [15,16].

Given the involvement of CRYPTOCHROME in dim light detection [17], we assayed the activity of a *cry* null mutant, *cry^01^*, under one of the bifurcation inducing regimes, LD^im^LD^im^ 5:7:5:7. As seen from the activity profiles (Fig 3A, top panel), *cry^01^* mutants did not display bifurcation. We next assayed activity under LD^im^LD^im^ 5:7:5:7 with 700 lux of bright light and 10 lux of dim light (Fig 3A, bottom panel). Thus, we increased the overall light intensity while maintaining the same contrast.

**Fig 3.**
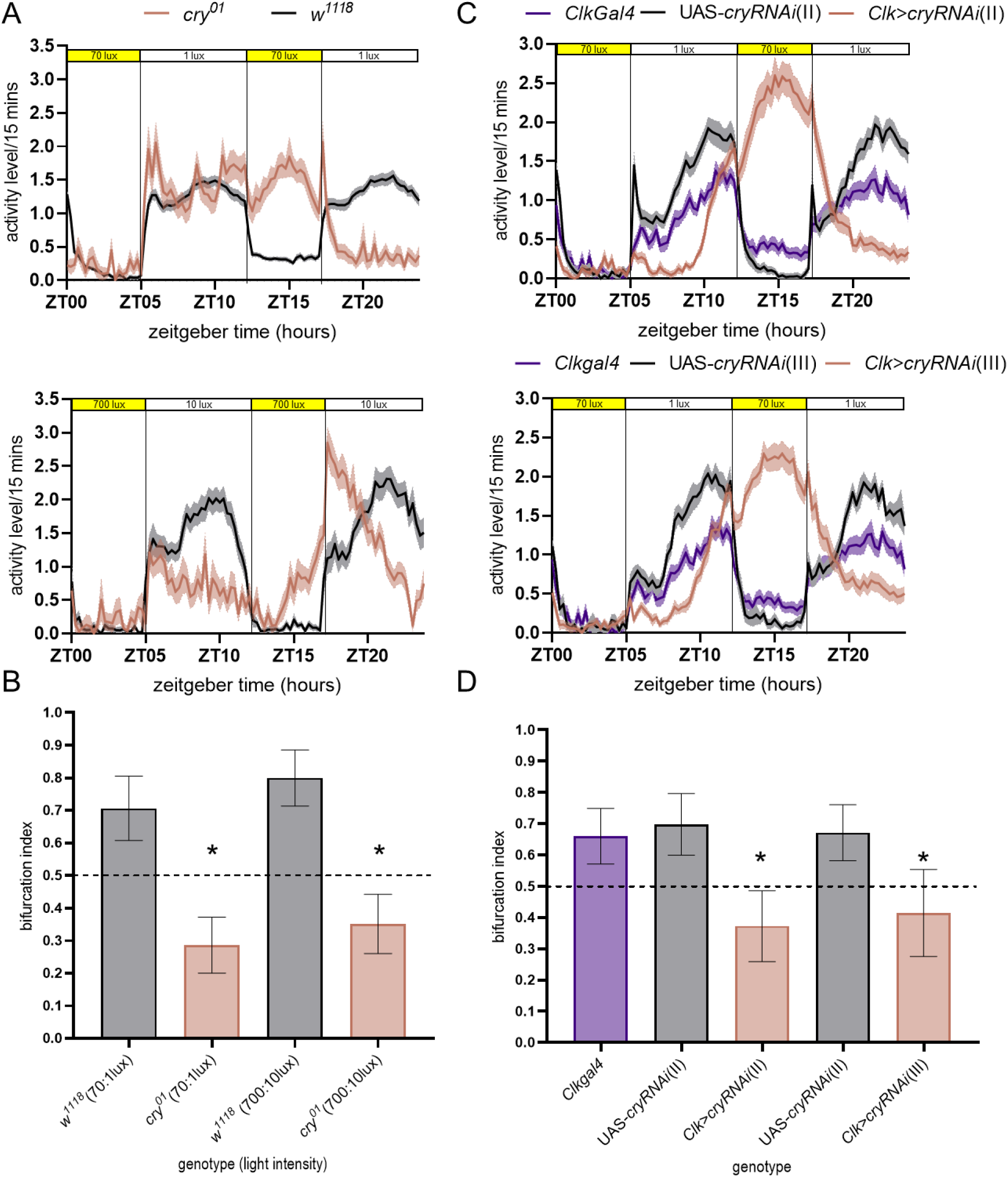
Role of CRYPTOCHROME in activity bifurcation. (A) Average activity profiles of *cry^01^*and its background control *w^1118^* under LD^im^LD^im^ 5:7:5:7 with the bright light of 70 lux, dim light of 1 lux (upper), and bright light of 700 lux with dim light of 10 lux (lower). Error bands are SEM. (B) Bifurcation indices of *cry* mutant and its background control, under the regimes with 70:1 lux of bright: dim light and 700: 10 lux of bright: dim light. The asterisks over the bars show a BI significantly lower than 0.5 by a one-sample *t*-test (*p* < 0.0001). For all strains, *n* > 20. Error bars are SD. (C) Activity profiles of flies with *cry* knocked down in the circadian clock neurons and their respective parental controls under LD^im^LD^im^ 5:7:5:7. Profiles of fly lines with two different *cryRNAi* constructs on the second and third chromosomes are depicted in the upper (VDRC-KK 103414) and lower (VDRC-GD 738) panels, respectively. (D) Bifurcation indices of experimental flies and their respective parental controls. Error bars are SD, and asterisks indicate BI significantly lower than 0.5 for the two experimental genotypes. BI was significantly higher than 0.5 for the parental controls by one-sample *t*-tests (*p* < 0.0001 for all genotypes except *clk>cryRNAi,* for which *p* = 0.0031). For each genotype *n* > 25.

These absolute light intensities fall within the detection range by HB eyelets and the compound eyes [15,16]. Even under the high light intensities, activity did not bifurcate in the mutants (Fig 3A, bottom), although the activity amplitude was higher. Further, we observed greater suppression of activity during the photophases under higher light intensity than regimes with 70 lux of bright light. The activity waveform of the *cryptochrome* mutants was quantified, and the bifurcation index was significantly lower than 0.5 (Fig 3B). These results indicate that this bona fide circadian photoreceptor is required for bifurcation.

CRY is also expressed in tissues other than the circadian circuit, like the compound eyes [18,19]. Therefore, using the RNAi approach, we knocked down *cry* expression specifically in the circadian clock neurons using the *Clock856GAL4* driver combined with UAS*-dicer* and assayed activity under LD^im^LD^im^ 5:7:5:7 (70 and 1 lux of bright and dim light, respectively). It is evident from the activity profiles that no bifurcation occurred in the flies with *cry* knocked down in the clock neurons (Fig 3C). Two independent transgenic constructs, one from VDRC-GD and one from VDRC-KK RNAi collections, inserted in chromosomes 3 and 2, respectively, yielded similar results (Figure 3C, upper and lower panels, respectively). The bifurcation indices for the experimental flies were significantly lower than 0.5, while parental controls (UAS controls for both RNAi constructs and *Clock856GAL4* with UAS-*dicer*) exhibited bifurcation indices significantly higher than 0.5 (Fig 3D). Flies with *cryptochrome* knocked down using *timA3GAL4* were also assayed under constant light for rhythmicity to verify the fly lines (S2 Fig) since flies lacking CRY are expected to be rhythmic under constant light, while control flies should be arrhythmic [20,21].

Thus, these experiments suggest that cell-autonomous CRY-mediated photoreception in circadian pacemaker neurons is necessary for dim light-restricted bifurcation of locomotor activity. This aligns with the observation that even light intensities within the detection range of compound eyes and HB eyelets (700:10 lux) did not result in bifurcation.

### Activity of circadian clock mutants bifurcates with altered waveform

To assess the contribution of circadian clock genes to the bifurcated waveform, we used null mutants of the components of the primary transcription-translation feedback loop in flies – *period*, *timeless*, *Clock* and *cycle*. Null mutants of *period* and *timeless* are known to show rhythms under synchronising conditions of LD 12:12 due to startle responses to light-dark transitions [20,22]. *Clock* and *cycle* null mutants also show startle responses, but their activity rhythm is weak [22,23]. A previous report showed that under LM 12:12 with moonlight intensity of 0.03 lux, clock mutants exhibit low-light induced activity, and the onset of nocturnal activity appears masked [24]. We, thus, expected startle responses and dim light-induced activity to be intact in clock mutants and aimed to assess the contribution of the circadian genes in regulating the waveform under LD^im^LD^im^ 5:7:5:7.

We exposed flies with *period*, *timeless*, *Clock* and *cycle* null mutations and their respective background controls to LD^im^LD^im^ 5:7:5:7 and, examined their profiles and quantified the bifurcation index (Fig 4). Startle responses to bright-dim transitions were observed in the mutant flies (Fig 4A-D). However, the overall activity waveform was altered compared to their control genotypes (Fig 4A-D). This change in the overall shape of the waveform suggests that clock genes contribute significantly to the timing of activity peaks and offsets under LD^im^LD^im^.

**Fig 4.**
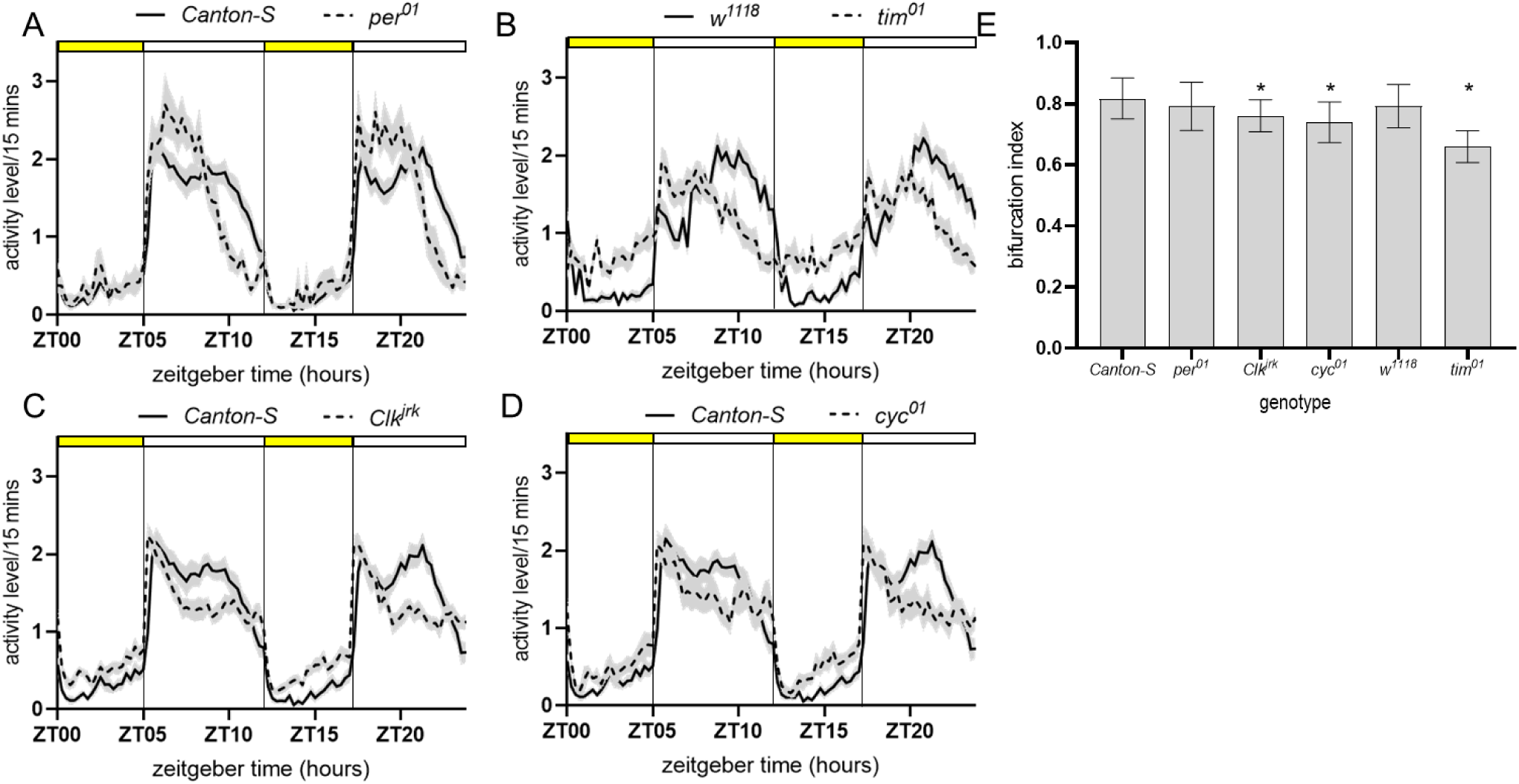
Role of circadian clock genes in the regulation of activity waveform. (A-D) Activity profiles with SEM error bands of the indicated clock gene mutants with their respective controls. (E) Bifurcation indices of all the genotypes. The values of each of the clock mutants were compared with the respective background controls. An asterisk indicates that the genotype had significantly lower BI than its control genotype. Error bars are SD. The values for *the period, Clock, and cycle* mutants and their background control were compared using a one-way ANOVA (*p* < 0.05, *n* > 12) followed by Tukey’s post hoc comparisons and the *timeless* mutant was compared to its control genotype using an unpaired *t*-test (*p* < 0.0001, *n* > 15). The bifurcation indices for all the six indicated genotypes were significantly greater than 0.5 by one-sample *t*-test, *p* < 0.001.

The above results suggest that the canonical components of the molecular circadian clock and its cell-autonomous photoreceptor play critical roles in determining the waveform under novel light regimes. This led us to investigate the contribution of the circadian clock circuit and its components in enabling such a distinct pattern of rhythmic behaviour.

### The evening cell group (LNds) governs the phases of both activity bouts

In the mouse model system, activity bifurcation was found to be correlated with antiphasic oscillations between the core and shell regions of the suprachiasmatic nucleus for the circadian proteins PER1 and BMAL1 (Watanabe et al., 2007; Yan et al., 2010). This suggests that the circadian pacemaker neuronal circuit undergoes reorganisation, and its components are likely decoupled due to entrainment to these novel light regimes. We aimed to utilise the genetic toolkit of *D. melanogaster* to understand the contributions of components of the pacemaker circuit in enabling this altered waveform of activity. Under LD 12:12, flies exhibit two bouts of activity centred around the dawn and dusk transitions. Previous work has shown that the morning and evening bouts of activity are strongly modulated by subsets of circadian neurons, often referred to as morning (M) and evening (E) cells, respectively [25,26]. The M group mainly comprises the large and small ventral lateral neurons (l-LNvs and s-LNvs) expressing the neuropeptide pigment dispersing factor (PDF). On the other hand, the canonical E cells primarily comprise the dorsal lateral neurons and a small ventral lateral neuron lacking the neuropeptide PDF (CRY^+ve^ LNds and 5^th^ s-LNv).

To examine the contribution of the M and E neurons to the waveform observed under novel light regimes, we overexpressed *DBT ^s^* in subsets of the canonical circadian pacemaker circuit. Using this approach, we could alter a parameter of the molecular clock, i.e. speed, within specific groups of neurons. As expected, both activity bouts were advanced under LD 12:12 in flies, with the allele being expressed in all the circadian neurons (S3 Fig). Only the morning bout of activity advanced when *DBT ^s^* was expressed in the PDF^+ve^ neurons, while only the evening bout advanced when *DBT ^s^* was expressed in the LNds, which are significant components of the evening group of neurons (S3 Fig)

Each genotype was then exposed to LD^im^LD^im^ 5:7:5:7. We observed that both activity bouts were advanced in flies with *DBT ^s^* expression in the circadian neurons compared to the control genotypes, indicating the regulation of the waveform by the circadian circuit. The effect was pronounced, especially for the peaks and the offsets of the two activity bouts. This was in line with our previous result, which indicated that the onset/dim light-induced activity is non-clock mediated. There were no differences in the activity waveform of the flies with only the morning cells expressing *DBT ^s^*. Interestingly, the expression of *DBT ^s^* only in the LNds/evening cells caused both bouts to advance. We used the centre of mass to estimate the mean phase of activity within each scotophase. The value would thus indicate the mean time around which the activity is centred within each scotophase. Expression of *DBT ^s^* in all the circadian cells (*Clk>DBT ^s^*) resulted in a significantly advanced mean phase of activity compared to its controls (UAS*-DBT ^s^* and *ClkGal4*) (Fig 5A). However, there was no difference in the mean phase of flies with a faster clock in M cells (*pdf>DBT ^s^*) relative to their parental controls (UAS*-DBT ^s^* and *pdfGal4*) (Fig 5B). In contrast, the expression of *DBT ^s^* only in E cells (*LNd>DBT ^s^*) was sufficient to significantly advance the mean phase of both activity bouts compared to its control genotypes (UAS*-DBT ^s^* and *LNdGal4*) (Fig 5C). Together, these results indicate that the circadian clocks in the evening cells of the pacemaker circuit modulate this unique behavioural pattern.

**Fig 5.**
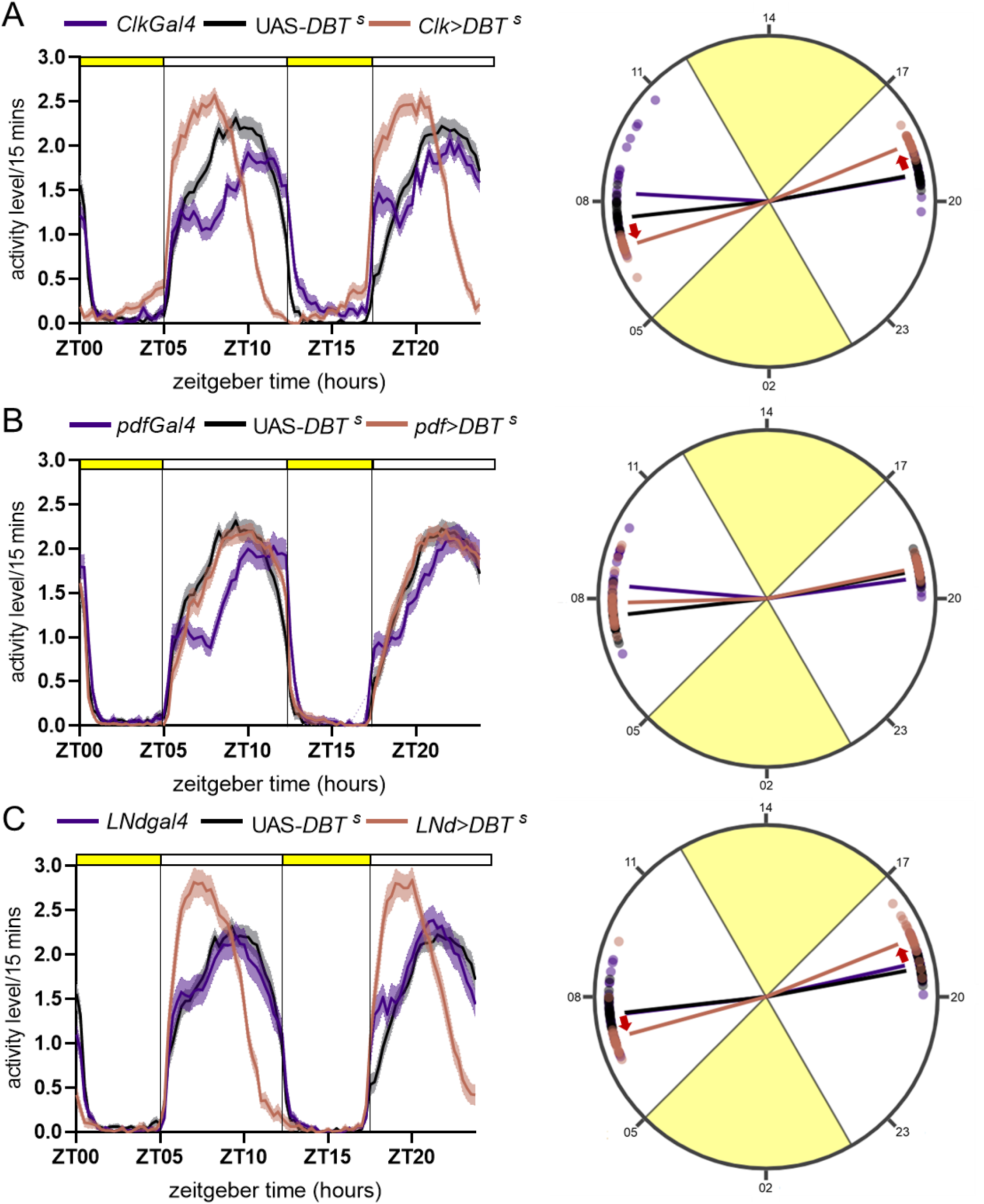
E cells govern the activity phases of the bifurcated activity bouts. On the left are activity profiles of flies with *DBT ^s^* expressed in circadian clock neurons or subsets of the circuit. Profiles have been averaged over days and then across individuals and are depicted with SEM error bands. Yellow and white bars on the top indicate bright and dim light, respectively. On the right are the mean phases of activity of each of the two bouts in scotophases (dim light). The phase estimated is the Centre of Mass (CoM), which considers the entire bout of activity during each scotophase. Phases of individual flies are circularly plotted with zeitgeber time on the circular axis, and the length of the vector indicates the consolidation of the mean phase values across individuals. For all genotypes *n* > 26. Red arrows in the circular plots indicate that the experimental genotype’s activity phase significantly differs from parental controls (Welch’s ANOVA, post hoc Dunett’s T3 multiple comparisons, *p* < 0.001). (A) Activity profiles and mean phases of flies with all circadian clock neurons expressing the *DBT ^s^*. (B) Activity profiles and phases of flies with only the PDF^+ve^ circadian neurons expressing the *DBT ^s^*. (C) Activity profiles and phases of flies with only the dorsolateral (LNds) circadian neurons expressing *DBT ^s^*.

In summary, under LD 12:12, the respective neuronal groups regulate morning and evening bouts. However, under LD^im^LD^im^, the evening neurons regulate the timing of both bouts of activity. Thus, we found evidence for circadian reorganisation under LD^im^LD^im^ compared to LD 12:12 using this approach.

## Discussion

The waveform of biological rhythms is altered in response to changes in the environmental cycles. Methods for waveform manipulation could be of tremendous use for practical purposes, such as easing the system to synchronise to challenging regimes like shift work or resynchronisation during jetlag or photoperiodic changes. The waveform of biological rhythms has also long served as a proxy to gauge the state of the underlying circuitry across model systems, including *D. melanogaster*. Analysing the waveform has also allowed the formation and testing of hypotheses regarding the wiring and coupling within the circadian circuit. The morning and evening oscillator model is a prime example. Here, we characterised Drosophila activity rhythm under novel light regimes as a means to systematically study the extent of plasticity of the activity waveform and its underlying circuitry.

We find that dim scotopic illumination (LD^im^LD^im^) induces and maintains a bifurcated activity pattern in *D. melanogaster*. Activity is consolidated in the dim light phases with little to no activity during the photophases. This indicates that the effect of scotopic illumination, leading to behavioural decoupling, is similar in mice and flies. Previous work with fruit flies shows that they prefer dimmer light intensities and exhibit increased nocturnal activity under dim nighttime illumination [12,27]. However, the use of novel light regimes revealed a phenomenon wherein flies exhibit activity exclusively in the dim-lit phases (Fig 1A). Furthermore, our results indicate that the duration of the scotophases is crucial to the induction of bifurcation. A light regime with a very short duration of the first scotophase did not induce activity bifurcation, suggesting that a threshold duration exists for behavioural decoupling to occur (Fig 2D). This is similar to the finding in rodents wherein specific durations of the photophases and scotophases induced activity bifurcation. While our study demonstrates that in flies, too, there are threshold durations for inducing bifurcation, we cannot yet pinpoint a value for that threshold duration. Nevertheless, our results reveal that certain factors contributing to activity bifurcation are conserved across flies and rodents. This bifurcated activity pattern, once adopted, is highly stable across cycles, indicating that particular light regimes can help ease the system to arrive at such an equilibrium. While specific combinations of the durations are critical for inducing activity bifurcation, the spectral quality of light could also be crucial to enabling such synchronisation. Another aspect would be to assess the impact on physiology upon exposure to such light regimes. Even though we did not conduct such a study, it is warranted since it can inform us about the practicality of using these regimes for waveform manipulation.

In mice, the bifurcated activity pattern was correlated with the reorganisation of the expression pattern of the circadian clock proteins in the suprachiasmatic nucleus of mice [8]. Using the genetic toolkit of the fly model system, we gained insights into the mechanistic components regulating activity bifurcation.

In both model systems, dim scotopic illumination was crucial for inducing activity bifurcation. Our study revealed that the dim light photoreceptor CRYPTOCHROME is necessary for this behaviour (Fig 3A). Additionally, we show that its function in the circadian neurons is essential for activity bifurcation (Fig 3C). Although not quantitatively comparable, it is noteworthy that when *cry* is absent in the brain and the compound eyes (*cry^01^*), the activity onset is tied to the onset of dim light; however, when it is absent only in the clock neurons, the activity is aligned to a photophase and appears like that under a long photoperiod. This observation hints that CRY may function differently in the pacemaker neurons versus in the compound eyes, where other photoreceptors are also present.

Thus, while compound eyes may contribute to the observed waveform, cell-autonomous CRY-mediated photoreception within the circadian pacemaker neurons is necessary for activity bifurcation (Fig 3C and D). Despite the presence of multiple photoreception pathways in flies, the role of a dedicated circadian photoreceptor in the pacemaker circuit is crucial for adopting such a distinct temporal niche. Hence, in mammals, it would be interesting to know if dim light detecting components in the rod cells or melanopsin in ipRGCs are responsible for the pattern observed.

A previous study assessing activity waveform under LM 12:12 demonstrated that moonlight-induced activity persisted even in circadian mutants [24]. Under LD^im^LD^im^ regimes, dim light-induced activity is intact in flies with mutations in the canonical clock genes. However, the activity waveform of these clock mutants is markedly different from their background controls (Fig 4), indicating that the clock regulates the phase of the activity peak and offset of the two bouts. These observations in circadian mutants suggest that while dim light-induced activity is independent of the clock, the timing of activity, particularly its peak and offset, is determined by canonical clock genes (Fig 4).

While these canonical clock genes are rhythmically expressed across the circadian pacemaker circuit, the M and E cells have distinct roles in determining the state of the circuit and, thus, the waveform. The behavioural decoupling observed in our study was also accompanied by an alteration in the dynamics of the circadian network compared to LD 12:12. Under this novel regime, we find that the E cells regulate the phasing of both activity bouts. Previous studies have demonstrated that the activity waveform of *Drosophila melanogaster* exhibits remarkable overlap with the state of the circuit. For instance, under a long photoperiod, the morning peak is reduced to a startle, and a prominent evening peak is observed in accordance with the E cells, determining the phasing of activity under these conditions [28]. Similarly, under LM 12:12, a prominent evening peak is observed and rescuing the clock only in the E neurons (*Mai* positive, *pdf* negative) by overexpressing *period* in a *period* null background is sufficient for wild-type like activity pattern under LM 12:12 [29]. The latter study also found that under LM 12:12, two groups within the E neurons were desynchronised in these flies. The 5^th^ sLNv and 1 of the CRY^+ve^ LNds formed one group, and two other CRY^+ve^ LNds formed the other group. Analyses of the connectomic data also indicate the presence of two groups within the canonical E cells, i.e. the 5th sLNv and 1 of the CRY^+ve^ LNds forming one group and two other CRY^+ve^ LNds forming another group [30,31]. Here, for the first time, we report that under the novel light regime LD^im^LD^im^, the phasing of two symmetrical bouts of activity is driven by evening neurons. We hypothesise that this regime has decoupled the E cellular group completely, with each group regulating one bout of activity. This warrants further investigation using cell-specific drivers such that only particular subsets of the evening neurons can be targeted separately to study their role in timing each of the two bouts of activity. Though challenging to execute, such an experiment may provide further insights into the reorganisation of the circadian network when exposed to novel light regimes.

In the past decade, these novel light regimes of LD^ark^LD^ark^ and LD^im^LD^im^ have been used to probe the extent of plasticity of rodent activity waveform [32]. To explore the generalizability of the findings in these studies, previous work from our laboratory used the fly model to characterise the activity waveform under LD^ark^LD^ark^ [11]. Interestingly, like rodents, flies also did not exhibit bifurcation under LD^ark^LD^ark^. In the current study, we find that dim scotopic illumination and threshold durations of the scotophases and photophases are common factors regulating activity bifurcation in both model systems. We also find that circadian organisation is altered compared to LD 12:12. Our study reveals that the evening cells can regulate the phases of two comparable bouts of activity. Thus, we uncover a hitherto unknown degree of plasticity of the circadian circuit using these novel light regimes. These regimes have, therefore, proved helpful in probing the plasticity of the circadian system across taxa and have led to insights at the behavioural and neuronal levels.

A combination of features of the light regime, like the ratio of light intensities and specific durations of the scotophases and photophases, can enable the system to synchronise to unconventional light regimes uniquely. However, the temporal niches of the organisms assayed under these novel light regimes are nocturnal (rodents) or crepuscular (*D. melanogaster*). In both cases, activity was found to be restricted to dim light. These circadian pacemakers and the underlying neuronal circuitry of these two model systems have been extensively investigated. There is evidence, at behavioural and circuit levels, for the existence of two oscillators and that they can decouple to a certain extent.

Hence, it is essential to conduct such studies across organisms (from different taxa with varying temporal niches and/or circuitry) to delineate the utility of such regimes for waveform manipulation.

Taken together, it is evident that certain factors inducing waveform bifurcation are conserved between flies and rodents. This opens up the possibility of designing light regimes to ease the activity waveform to arrive at such an equilibrium. Thus, such studies can result in an understanding of factors and principles governing waveform plasticity and help arrive at a means to manipulate activity waveform for practical purposes like shift work.

## Materials and methods

### Fly strains

The strains used in this study include outbred populations of *Drosophila melanogaster* (fly populations referred to as *control* [11] and inbred strains of *Canton-S* and *w^1118^*. The mutant and transgenic lines used are as follows:

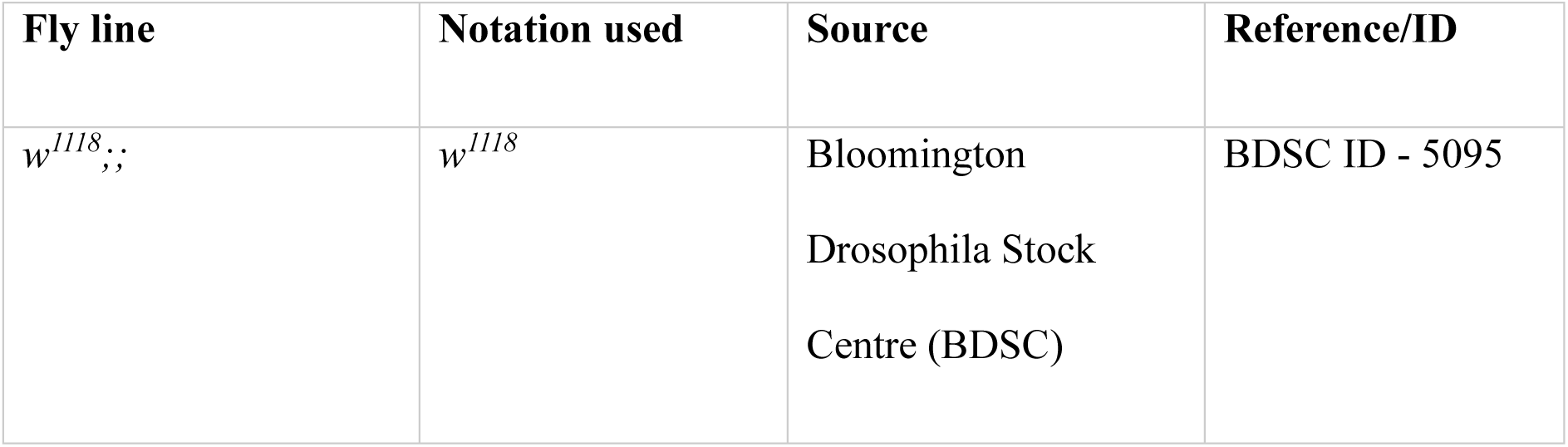
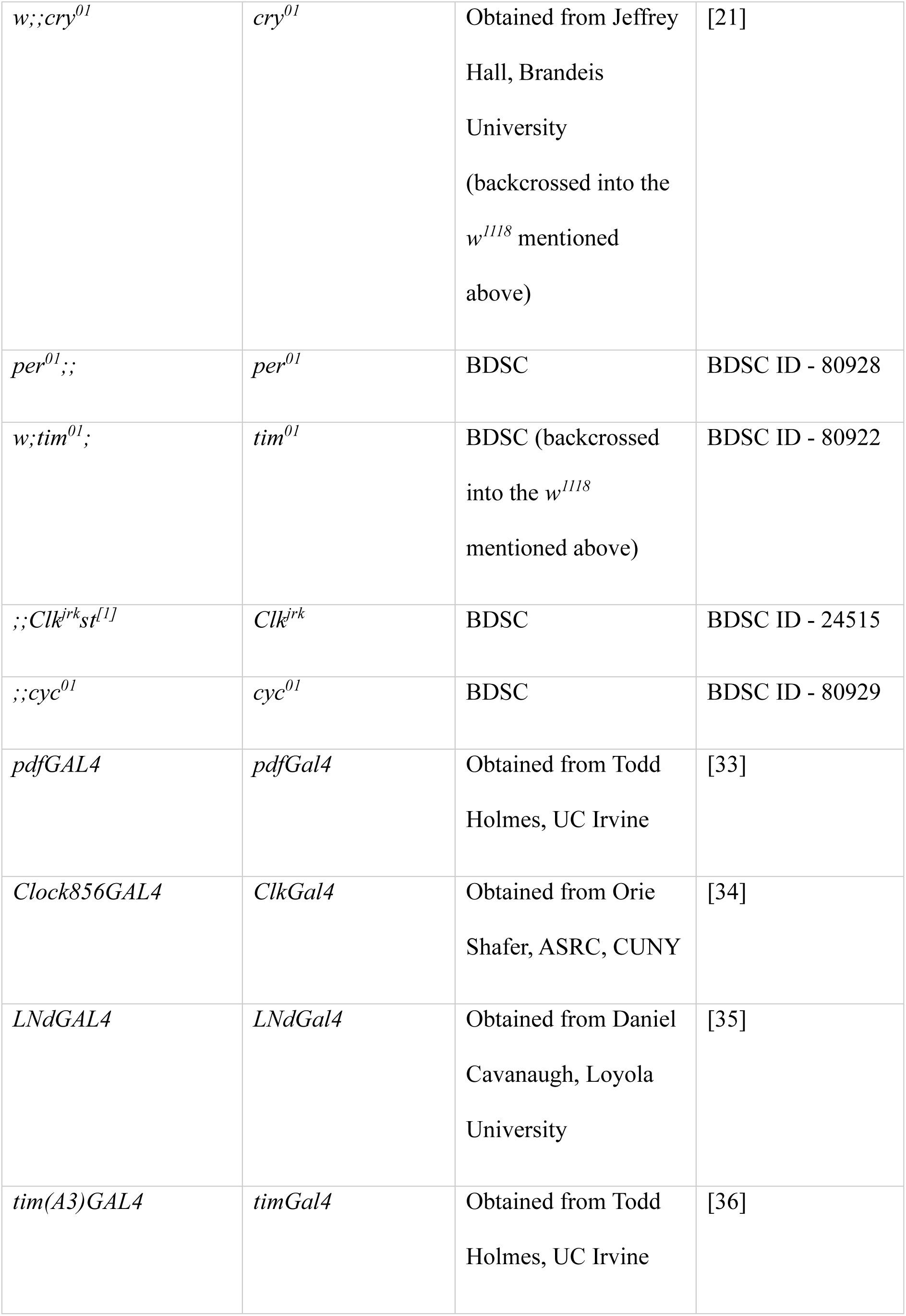
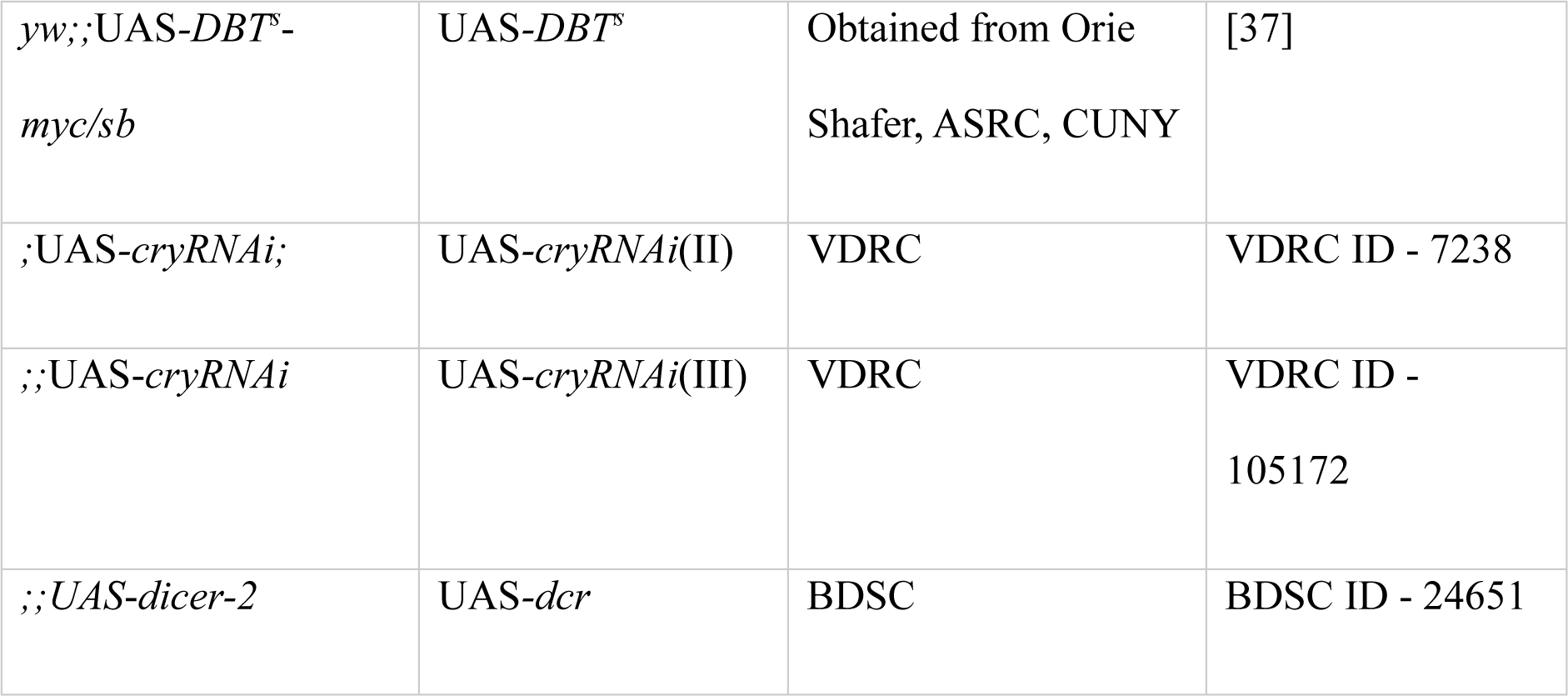

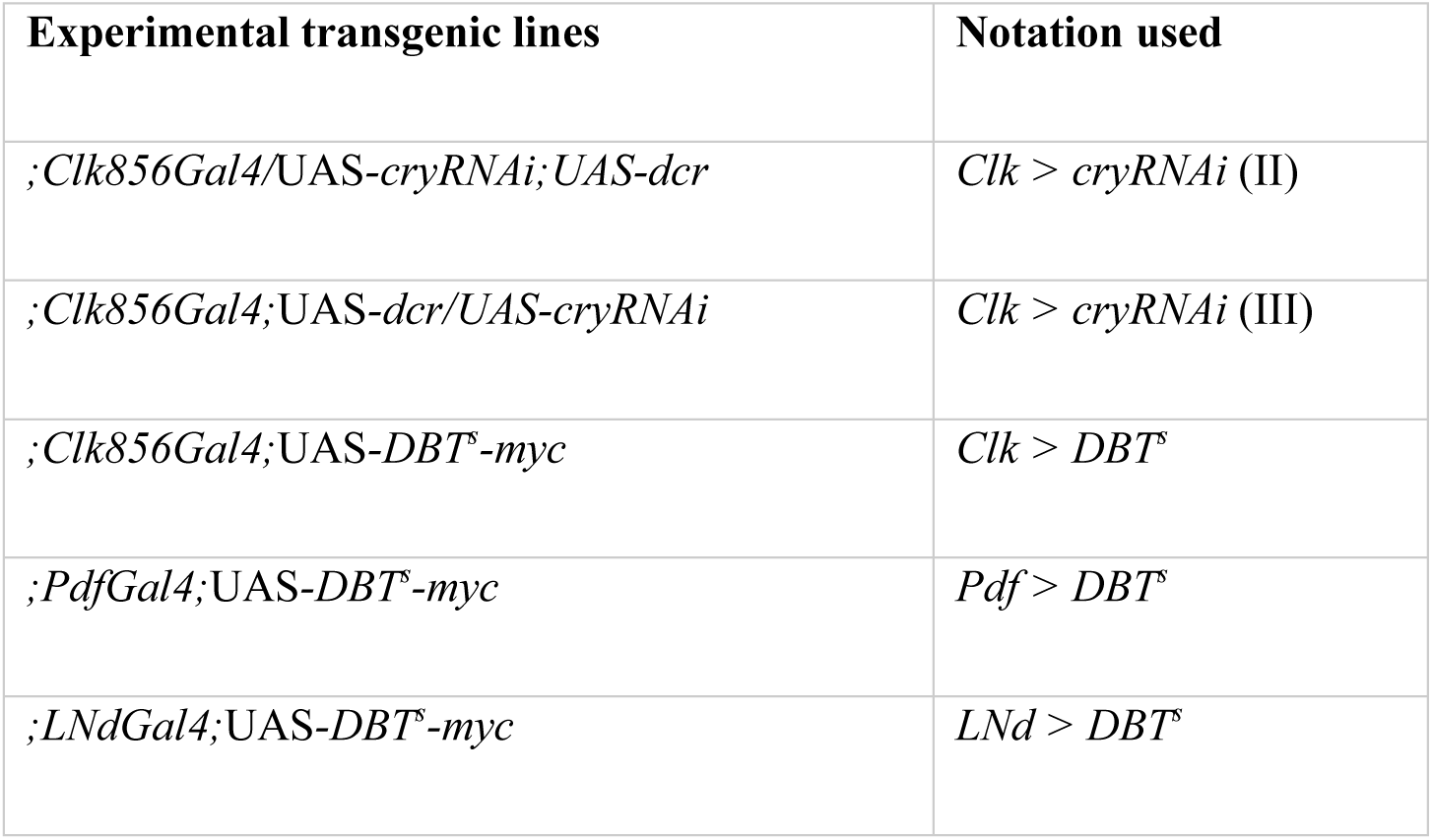

| Experimental transgenic lines | Notation used |
| --- | --- |
| *;Clk856Gal4/UAS-cryRNAi;UAS-dcr* | *Clk > cryRNAi* (II) |
| *;Clk856Gal4;UAS-dcr/UAS-cryRNAi* | *Clk > cryRNAi* (III) |
| *;Clk856Gal4;UAS-DBT<sup>s</sup>-myc* | *Clk > DBT<sup>s</sup>* |
| *;PdfGal4;UAS-DBT<sup>s</sup>-myc* | *Pdf > DBT<sup>s</sup>* |
| *;LNdGal4;UAS-DBT<sup>s</sup>-myc* | *LNd > DBT<sup>s</sup>* |

### Locomotor rhythm recording

3-5 days old virgin male flies were housed in locomotor tubes with corn food medium and placed in Drosophila Activity Monitors (DAM system by Trikinetics) to measure their activity counts. Assays were carried out in light-proof metal boxes placed in cubicles with an ambient temperature of 25°C ± 0.5°C. Light intensity was adjusted using a combination of 210 and 298 neutral-density Lee filters over white LED strips fitted inside the boxes. Light intensities were adjusted using a LI-COR light meter.

Flies were maintained under LD 12:12, 70 lux: 0 lux cycles, then shifted to LD^im^LD^im^ regimes. The first bright light/70 lux phase is referred to as photophase 1 (PP1), and its onset coincides with that of the photophase of the previous LD cycle. The first scotophase with dim light is referred to as scotophase 1 (SP1), and the following bright light and dim light phases are photophase 2 (PP2) and scotophase 2 (SP2), respectively. Four regimes have been used, with varying durations of each photophase or scotophase. In each case, the transfer from LD 12:12 to LD^im^LD^im^ was such that the start of PP1 coincided with the start of the 12-hour photophase of the previous day. Thus, the start of the first photophase is regarded as zeitgeber time 00 or ZT00 in this case.

### Data analyses

The bifurcation symmetry index has been used previously to quantify the bifurcated activity patterns observed in rodents [5]. Since we, too, observed activity restricted to the scotophases, we have used the same measure to assess the degree of symmetry in activity division between successive scotophases. The total number of activity counts in each scotophase (SP1, SP2) and the daily total activity was calculated for each cycle. The activity counts in the lesser-active scotophase are doubled to provide an objective measure of bifurcation ranging from 0 to 1. If activity is perfectly bifurcated between the successive scotophases, the BI = 1. Differences in activity level between the scotophases or any activity during the photophases will lower this index. Even if the activity is concentrated in one of the scotophases, the index will approach 0.

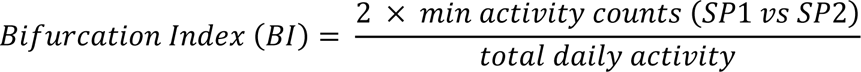

The circular mean phase of activity within the scotophases was also quantified for certain genotypes to measure the centrality of activity on the time axis [38]. It considers and quantifies the overall activity waveform.

All the activity profiles and bar graphs have been plotted using GraphPad Prism v.8. Statistical tests reported have also been performed using GraphPad Prism v.8.

## Supporting information

**S1 Fig.**
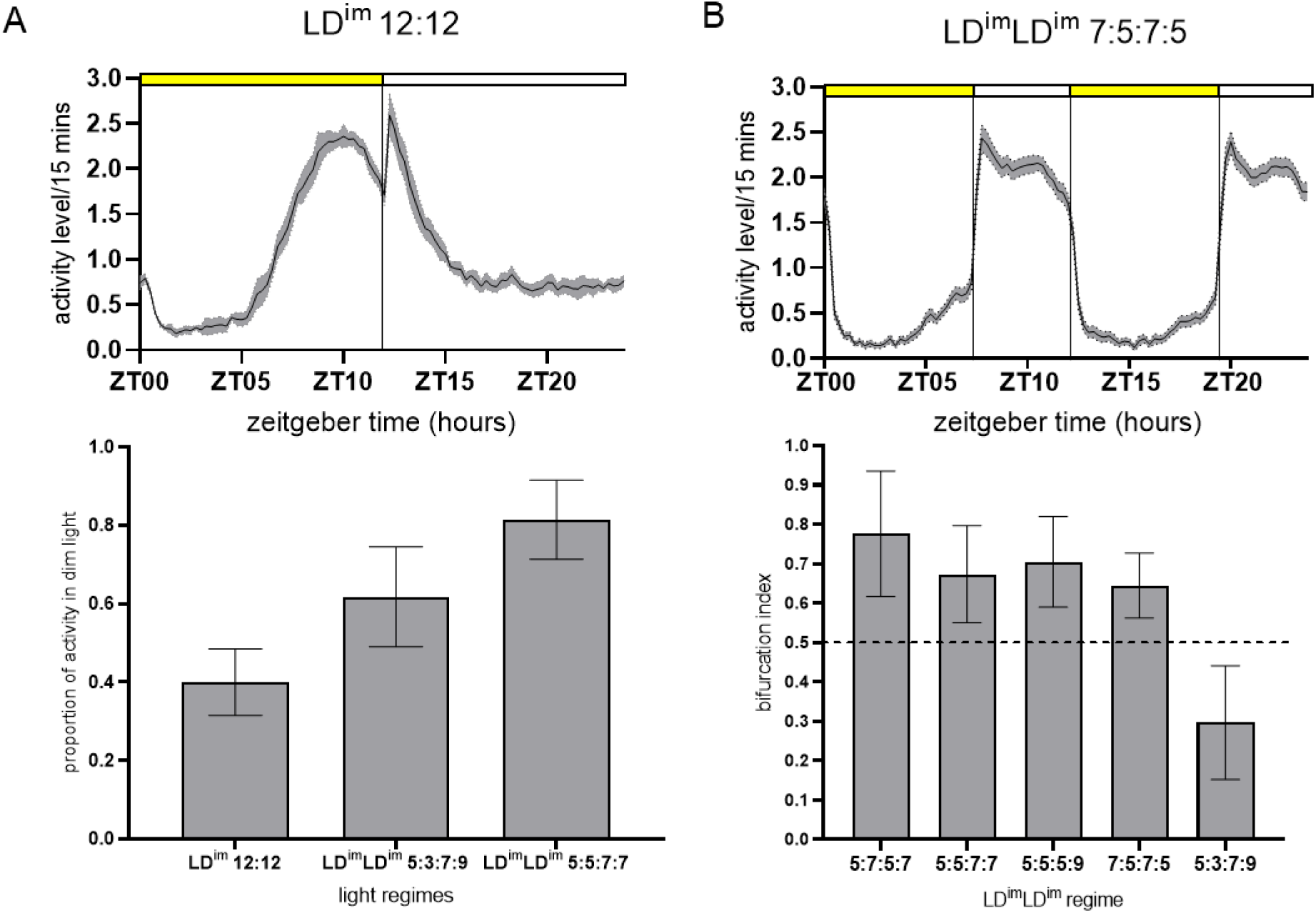
Quantification of activity distribution under light: dim regimes. (A) Upper panel: activity profile of flies under LD^im^ 12:12 with bright light of 70 lux and dim light of 1 lux. Lower panel: proportion of activity exhibited by flies during dim light, plotted with LD^im^LD^im^ regimes having the same total bright and dim light duration. Under bifurcation inducing regimes, activity is largely restricted to the dim light, unlike the other two regimes. Error bars are SD. (B) Upper panel: activity profile of GC flies under LD^im^LD^im^ 7:5:7:5 with bright light of 70 lux and dim light of 1 lux. Lower panel: bifurcation indices of flies under all the LD^im^LD^im^ regimes. Error bars are SD. For LD^im^LD^im^ 7:5:7:5, the bifurcation index was significantly greater than 0.5 by a one-sample *t*-test (*n* > 50, *p* < 0.0001).

**S2 Fig.**
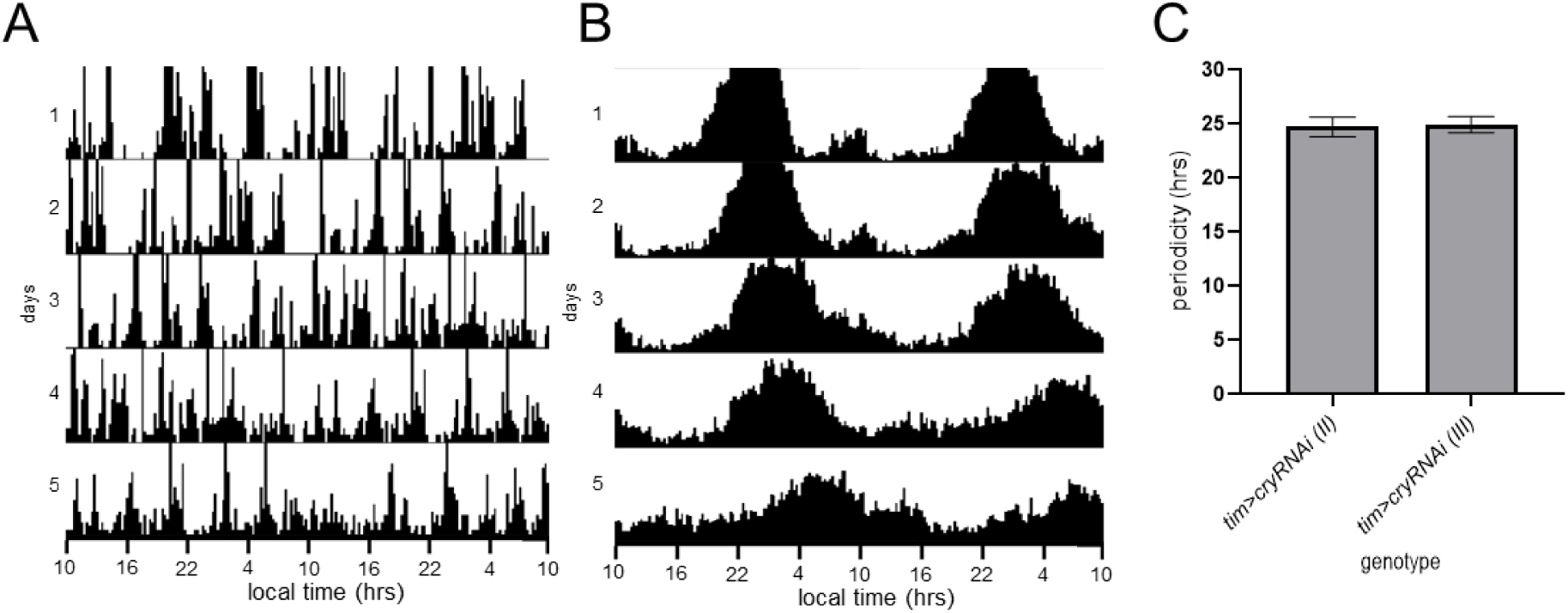
Phenotyping of flies with *cryptochrome* knocked down to verify *cryRNAi* fly lines. (A) Representative actogram of UAS control flies with the UAS*-cryRNAi* on the third chromosome under 1 lux constant light. For both RNAi constructs, UAS*-cryRNAi (II)* and UAS*-cryRNAi(III)*, > 80% of flies were arrhythmic. (B) Batch actogram of flies with *timGal4* driving the expression of UAS*-cryRNAi (III)*. All the individuals were rhythmic under constant 1 lux. Similarly, all the individuals with *timGal4* driving the expression of *UAS-cryRNAi (III)* were also rhythmic. (C) Average period value of the genotypes with *cryptochrome* knocked down using the *timGal4* driver under constant conditions with 1 lux light intensity. Error bars are SD, *n* > 19.

**S3 Fig.**
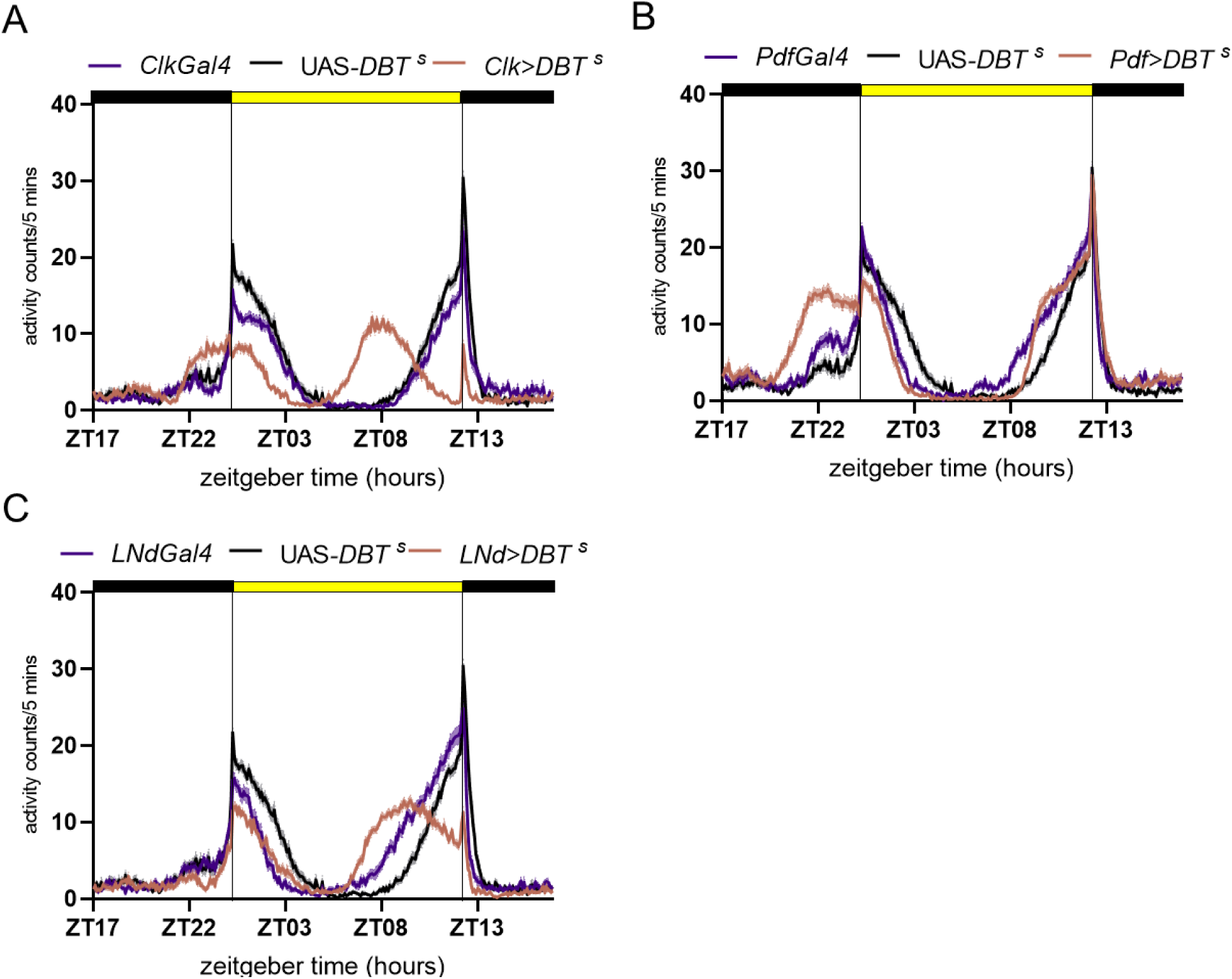
Activity profiles of flies with *DBT ^s^* overexpression using three driver lines under LD 12:12. As evident from the activity profiles, both activity bouts advance when *DBT ^s^* is overexpressed in all the circadian neurons. On the other hand, only the morning bout is advanced when *pdfgal4* is used, while only the evening bout advances when *LNdgal4* is driving the expression of *DBT ^s^*. The yellow and black bars indicate day and night durations, respectively.

**S1 Table.**
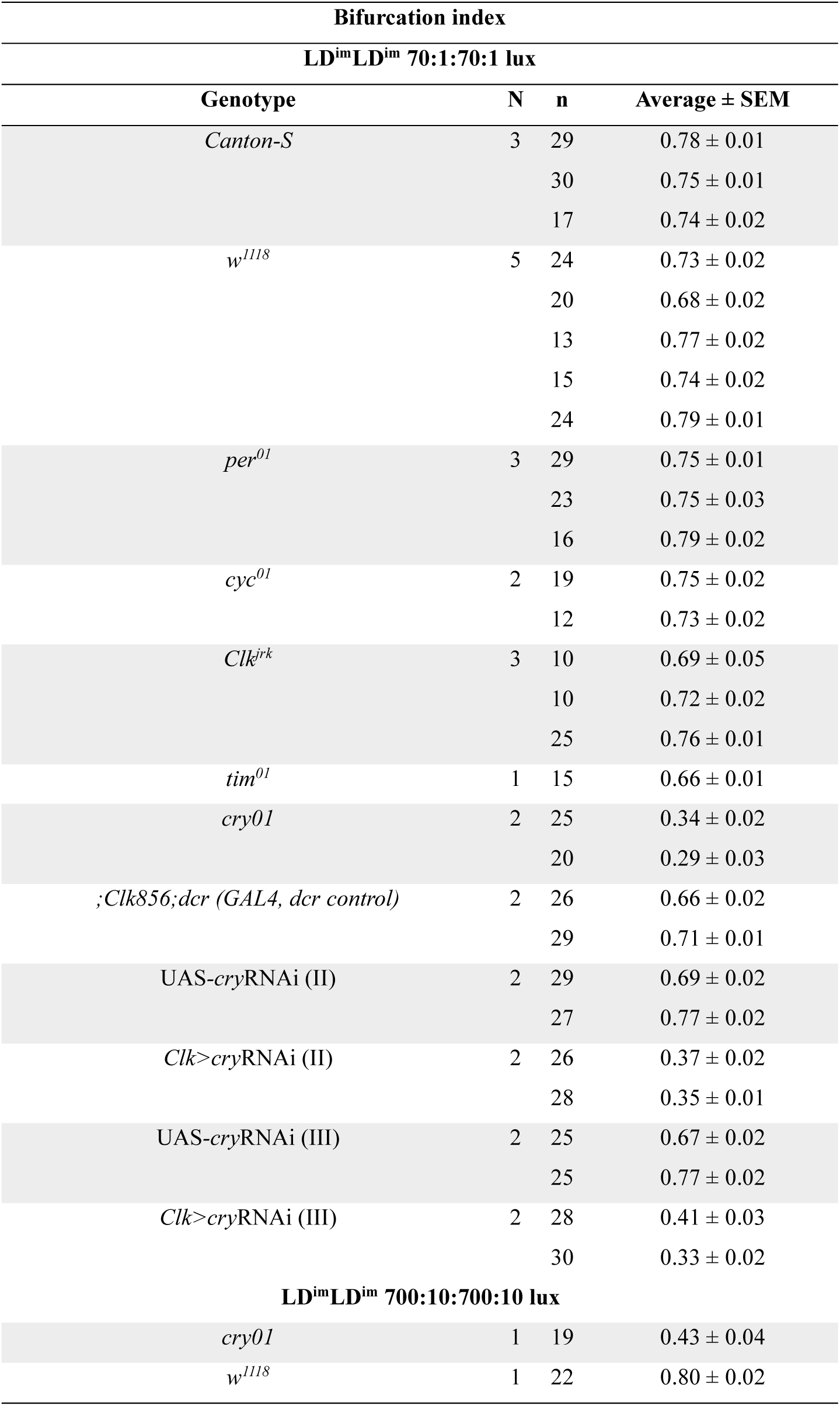
Bifurcation index under LD^im^LD^im^ light regime. Average bifurcation index (± SEM) of all the assayed genotypes with the number of experimental replicates and number of flies under LD^im^LD^im^ under the mentioned light intensities.

**S2 Table.**
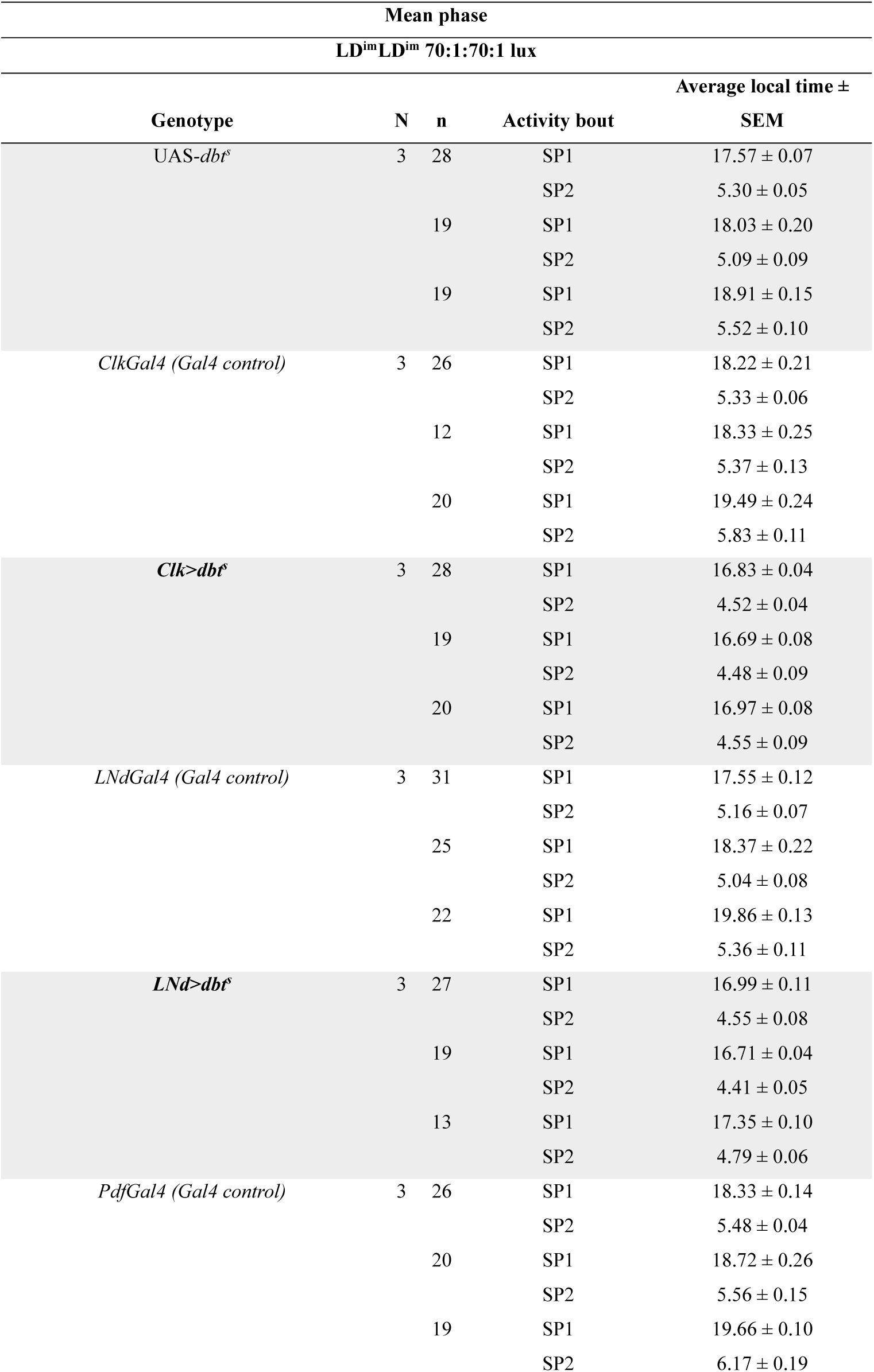
Mean phase values of flies with *dbt^s^* expression in circadian neurons and their respective controls. The table above lists the transgenic genotypes, number of replicate experiments, number of flies and the average mean phase in local time (± SEM) of each activity bout under LD^im^LD^im^. Both the bouts of activity were significantly advanced compared to their respective parental controls for the genotypes in bold.

## Notes

### Competing Interest Statement

The authors have declared no competing interest.

## References

1. Redlin U. Neural basis and biological function of masking by light in mammals: suppression of melatonin and locomotor activity. Chronobiol Int. 2001 Sep;18(5):737–58.

2. Sheppard AD, Hirsch HVB, Possidente B. Novel masking effects of light are revealed in Drosophila by skeleton photoperiods. Biological Rhythm Research. 2015 Mar 4;46(2):275–85.

3. Singh VJ, Potdar S, Sheeba V. Effects of Food Availability Cycles on Phase and Period of Activity-rest Rhythm in Drosophila melanogaster. J Biol Rhythms. 2022 Oct 1;37(5):528–44.

4. Evans JA, Elliott JA, Gorman MR. Circadian entrainment and phase resetting differ markedly under dimly illuminated versus completely dark nights. Behavioural Brain Research. 2005 Jul;162(1):116–26.

5. Harrison EM, Walbeek TJ, Sun J, Johnson J, Poonawala Q, Gorman MR. Extraordinary behavioral entrainment following circadian rhythm bifurcation in mice. Sci Rep. 2016 Dec 8;6(1):38479.

6. Gorman MR, Elliott JA. Dim nocturnal illumination alters coupling of circadian pacemakers in Siberian hamsters, Phodopus sungorus. J Comp Physiol A [Internet]. 2004 Aug [cited 2023 Apr 14];190(8). Available from: http://link.springer.com/10.1007/s00359-004-0522-7

7. Yan L, Silver R, Gorman M. Reorganization of Suprachiasmatic Nucleus Networks under 24-h LDLD Conditions. J Biol Rhythms. 2010 Feb;25(1):19–27.

8. Watanabe T, Naito E, Nakao N, Tei H, Yoshimura T, Ebihara S. Bimodal Clock Gene Expression in Mouse Suprachiasmatic Nucleus and Peripheral Tissues Under a 7-Hour Light and 5-Hour Dark Schedule. J Biol Rhythms. 2007 Feb;22(1):58–68.

9. Evans JA, Elliott JA, Gorman MR. Dynamic interactions between coupled oscillators within the hamster circadian pacemaker. Behavioral Neuroscience. 2010;124(1):87–96.

10. Gorman MR, Steele NA. Phase Angle Difference Alters Coupling Relations of Functionally Distinct Circadian Oscillators Revealed by Rhythm Splitting. J Biol Rhythms. 2006 Jun;21(3):195–205.

11. Abhilash L, Ramakrishnan A, Priya S, Sheeba V. Waveform Plasticity under Entrainment to 12-h *T* -cycles in *Drosophila melanogaster* : Behavior, Neuronal Network, and Evolution. J Biol Rhythms. 2020 Apr;35(2):145–57.

12. Bachleitner W, Kempinger L, Wülbeck C, Rieger D, Helfrich-Förster C. Moonlight shifts the endogenous clock of *Drosophila melanogaster*. Proc Natl Acad Sci USA. 2007 Feb 27;104(9):3538–43.

13. Shafer OT, Levine JD, Truman JW, Hall JC. Flies by Night: Effects of Changing Day Length on Drosophila’s Circadian Clock. Current Biology. 2004 Mar 9;14(5):424–32.

14. Sun J, Joye DAM, Farkas AH, Gorman MR. Photoperiodic Requirements for Induction and Maintenance of Rhythm Bifurcation and Extraordinary Entrainment in Male Mice. Clocks & Sleep. 2019 Jul 4;1(3):290–305.

15. Helfrich-Förster C. Light input pathways to the circadian clock of insects with an emphasis on the fruit fly Drosophila melanogaster. J Comp Physiol A. 2020 Mar 1;206(2):259–72.

16. Schlichting M, Menegazzi P, Rosbash M, Helfrich-Förster C. A Distinct Visual Pathway Mediates High-Intensity Light Adaptation of the Circadian Clock in Drosophila. J Neurosci. 2019 Feb 27;39(9):1621–30.

17. Vinayak P, Coupar J, Hughes SE, Fozdar P, Kilby J, Garren E, et al. Exquisite Light Sensitivity of Drosophila melanogaster Cryptochrome. PLOS Genetics. 2013 Jul 18;9(7):e1003615.

18. Mazzotta G, Rossi A, Leonardi E, Mason M, Bertolucci C, Caccin L, et al. Fly cryptochrome and the visual system. Proceedings of the National Academy of Sciences. 2013 Apr 9;110(15):6163–8.

19. Yoshii T, Todo T, Wülbeck C, Stanewsky R, Helfrich-Förster C. Cryptochrome is present in the compound eyes and a subset of Drosophila’s clock neurons. Journal of Comparative Neurology. 2008;508(6):952–66.

20. Collins BH, Dissel S, Gaten E, Rosato E, Kyriacou CP. Disruption of Cryptochrome partially restores circadian rhythmicity to the arrhythmic period mutant of Drosophila. Proceedings of the National Academy of Sciences. 2005 Dec 27;102(52):19021–6.

21. Dolezelova E, Dolezel D, Hall JC. Rhythm Defects Caused by Newly Engineered Null Mutations in Drosophila’s cryptochrome Gene. Genetics. 2007 Sep 1;177(1):329–45.

22. Allada R, White NE, So WV, Hall JC, Rosbash M. A Mutant *Drosophila* Homolog of Mammalian *Clock* Disrupts Circadian Rhythms and Transcription of *period* and *timeless*. Cell. 1998 May 29;93(5):791–804.

23. Rutila JE, Suri V, Le M, So WV, Rosbash M, Hall JC. CYCLE Is a Second bHLH-PAS Clock Protein Essential for Circadian Rhythmicity and Transcription of *Drosophila period* and *timeless*. Cell. 1998 May 29;93(5):805–14.

24. Kempinger L, Dittmann R, Rieger D, Helfrich-Förster C. The Nocturnal Activity of Fruit Flies Exposed to Artificial Moonlight Is Partly Caused by Direct Light Effects on the Activity Level That Bypass the Endogenous Clock. Chronobiology International. 2009 Jan;26(2):151–66.

25. Grima B, Chélot E, Xia R, Rouyer F. Morning and evening peaks of activity rely on different clock neurons of the Drosophila brain. Nature. 2004 Oct;431(7010):869–73.

26. Stoleru D, Peng Y, Agosto J, Rosbash M. Coupled oscillators control morning and evening locomotor behaviour of Drosophila. Nature. 2004 Oct;431(7010):862–8.

27. Rieger D, Fraunholz C, Popp J, Bichler D, Dittmann R, Helfrich-Förster C. The Fruit Fly *Drosophila melanogaster* Favors Dim Light and Times Its Activity Peaks to Early Dawn and Late Dusk. J Biol Rhythms. 2007 Oct;22(5):387–99.

28. Stoleru D, Nawathean P, Fernández M de la P, Menet JS, Ceriani MF, Rosbash M. The Drosophila Circadian Network Is a Seasonal Timer. Cell. 2007 Apr 6;129(1):207–19.

29. Rieger D, Wülbeck C, Rouyer F, Helfrich-Förster C. Period Gene Expression in Four Neurons Is Sufficient for Rhythmic Activity of Drosophila melanogaster under Dim Light Conditions. J Biol Rhythms. 2009 Aug 1;24(4):271–82.

30. Scheffer LK, Xu CS, Januszewski M, Lu Z, Takemura S ya, Hayworth KJ, et al. A connectome and analysis of the adult Drosophila central brain. Marder E, Eisen MB, Pipkin J, Doe CQ, editors. eLife. 2020 Sep 3;9:e57443.

31. Shafer OT, Gutierrez GJ, Li K, Mildenhall A, Spira D, Marty J, et al. Connectomic analysis of the Drosophila lateral neuron clock cells reveals the synaptic basis of functional pacemaker classes. Desplan C, Helfrich-Förster C, editors. eLife. 2022 Jun 29;11:e79139.

32. Gorman MR, Harrison EM, Evans JA. Circadian Waveform and Its Significance for Clock Organization and Plasticity. In: Kumar V, editor. Biological Timekeeping: Clocks, Rhythms and Behaviour [Internet]. New Delhi: Springer India; 2017 [cited 2024 Mar 1]. p. 59–79. Available from: 10.1007/978-81-322-3688-7_4

33. Renn SCP, Park JH, Rosbash M, Hall JC, Taghert PH. A pdf Neuropeptide Gene Mutation and Ablation of PDF Neurons Each Cause Severe Abnormalities of Behavioral Circadian Rhythms in Drosophila. Cell. 1999 Dec;99(7):791–802.

34. Gummadova JO, Coutts GA, Glossop NRJ. Analysis of the Drosophila Clock Promoter Reveals Heterogeneity in Expression between Subgroups of Central Oscillator Cells and Identifies a Novel Enhancer Region. J Biol Rhythms. 2009 Oct 1;24(5):353–67.

35. Bulthuis N, Spontak KR, Kleeman B, Cavanaugh DJ. Neuronal Activity in Non-LNv Clock Cells Is Required to Produce Free-Running Rest:Activity Rhythms in Drosophila. J Biol Rhythms. 2019 Jun 1;34(3):249–71.

36. Kaneko M, Hall JC. Neuroanatomy of cells expressing clock genes in Drosophila: Transgenic manipulation of the period and timeless genes to mark the perikarya of circadian pacemaker neurons and their projections. Journal of Comparative Neurology. 2000;422(1):66–94.

37. Muskus MJ, Preuss F, Fan JY, Bjes ES, Price JL. Drosophila DBT Lacking Protein Kinase Activity Produces Long-Period and Arrhythmic Circadian Behavioral and Molecular Rhythms. Molecular and Cellular Biology. 2007 Dec 1;27(23):8049–64.

38. Batschelet E. Circular Statistics in Biology. Academic Press; 1981. 371 p.

